# The diversity of quinoa morphological traits and seed metabolic composition

**DOI:** 10.1101/2021.07.17.452781

**Authors:** Iman Tabatabaei, Saleh Alseekh, Mohammad Shahid, Ewa Leniak, Mateusz Wagner, Henda Mahmoudi, Sumitha Thushar, Alisdair R. Fernie, Kevin M. Murphy, Sandra M. Schmöckel, Mark Tester, Bernd Mueller-Roeber, Aleksandra Skirycz, Salma Balazadeh

## Abstract

Quinoa (*Chenopodium quinoa* Willd.) is an herbaceous annual crop of the amaranth family (Amaranthaceae). It is increasingly cultivated for its nutritious grains, which are rich in protein and essential amino acids, lipids, and minerals. Quinoa exhibits a high tolerance towards various abiotic stresses including drought and salinity, which supports its agricultural cultivation under climate change conditions. The use of quinoa grains is compromised by anti-nutritional saponins, a terpenoid class of secondary metabolites deposited in the seed coat; their removal before consumption requires extensive washing, an economically and environmentally unfavorable process; or their accumulation can be reduced through breeding. In this study, we analyzed the seed metabolomes, including amino acids, fatty acids, and saponins, from 471 quinoa cultivars, including two related species, by liquid chromatography – mass spectrometry. Additionally, we determined a large number of agronomic traits including biomass, flowering time, and seed yield. The results revealed considerable diversity between genotypes and provide a knowledge base for future breeding or genome editing of quinoa.

|  |  |
| --- | --- |
| Design type | Field-grown quinoa genotypes |
| Measurement Type(s) | Agronomic traits and seed metabolites |
| Technology Type(s) | Agronomic trait analysis and liquid chromatography –mass spectrometry (LC-MS) |
| Factor Type(s) | Genotypes |
| Sample Characteristic(s) | Quinoa (*Chenopodium quinoa* Willd.), agricultural traits, composition of the seed metabolome |

## Background & Summary

Quinoa (*Chenopodium quinoa* Willd.) is increasingly attracting global attention because of its unusually high grain nutritional value including high protein content, the composition and quantity of lipids, a good balance of essential amino acids, as well as isoflavones and interesting antioxidant functional properties [1–3]. Quinoa was first domesticated in the Lake Titicaca basin about 7,000 years ago, from where it spread to other regions in South America and the world [4].

An agriculturally important asset of quinoa is its remarkable ability to adapt to diverse agroecological zones, which allows growth in hot dry deserts and in tropical areas with up to 88% relative humidity, from −8°C to 40°C [5], and from sea level to 4,000 m high mountainous regions. Its adaptability to sodic and alkaline soils is also remarkable allowing cultivation from pH 4.5 to 9.0 [6]. Quinoa is a highly drought tolerant crop that fares well in regions of below 200 mm yearly rainfall ([7] and references therein). It is tolerant against high salinity and considered a facultative halophyte [8–10]. In 2013, the Food and Agricultural Organization (FAO) declared the ′International Year of Quinoa′ in recognition of the capacity of the crop to help mitigate hunger and malnutrition in food-insecure countries, and in recognition of the ancestral efforts of the Andean people to preserve quinoa as a crop (http://www.fao.org/quinoa-2013/en/).

Although quinoa grains have an exceptional nutritional value, the seed coat typically contains bitter-tasting and potentially anti-nutritional saponins [11]. Therefore, quinoa seeds require substantial processing (water-extensive washing) to remove saponins before consumption. Reduction of saponins has been a breeding target and in the future may also be achieved with biotechnological methods, such as genome editing. The quinoa saponins occur predominantly in the form of triterpenoid glycosides [12–14]. Their large structural diversity renders analyses non-trivial [15].

The biological functions of saponins in quinoa remain to be investigated. Saponins may play a role in seed germination, and in deterring birds or fungal infections (reviewed in [16]). Evidence indicates that not only the total amount of saponins is regulated (e.g., by the bHLH transcription factor CqTSARL1) [17], but also the saponin profile [17]. However, to date, seed saponin profiles of only few quinoa genotypes have been determined [17]. As some saponins may even be beneficial to human health [18], the diversity in saponin composition poses a great resource for breeding new and more healthy quinoa cultivars.

Our study reports the variability of the metabolome of mature quinoa seeds of a large number of genotypes (471 in total; **Supplementary Dataset 1**). Additionally, we determined agronomic traits such as plant height, total biomass, panicle density, days to flowering, and seed weight (**Fig. 1** and **Supplementary Table 1**). The experimental pipeline employed for liquid chromatography – mass spectrometry (LC-MS)-based metabolome analysis of seeds is represented in Figure 2. Metabolites were annotated using a library of authentic reference compounds, and in-source fragmentation patterns, and the data are reported in compliance with established standards [19] (**Supplementary Table 2** and MetaboLights database, MTBLS2382). We detected and quantified 400 seed metabolites representing diverse chemical classes: 37 triterpenoid saponins, 14 flavonoids, 15 amino acids, 117 dipeptides, 126 lipids, and 91 other metabolites. To explore the variation between genotypes, principal component analysis (PCA) and a hierarchical cluster (HCA) heatmap were established on metabolic and phenotypic traits (**Fig. 3a, b**). The heatmap revealed considerable differences in metabolite abundances across genotypes (**Fig. 3a**), which was confirmed by PCA (**Fig. 3b**), in which the first and the second components explained 41.1% and 19.9% respectively, of saponin variance. PCA analysis identified 21 genotypes, mostly originating from Peru/Latin America, whose position in the PCA plot largely correlates with their geographical origin (**Supplementary Table 3**). Finally, we investigated correlations between and within different metabolite classes and phenotypic traits (**Fig. 3c**). This showed that many metabolites are highly associated within the network. However, no significant strong correlations were found between saponin content and morphological traits, indicating that genotypes with low saponin content can be selected in the future by breeding or genome editing without an impact on yield-related traits.

**Figure 1.**
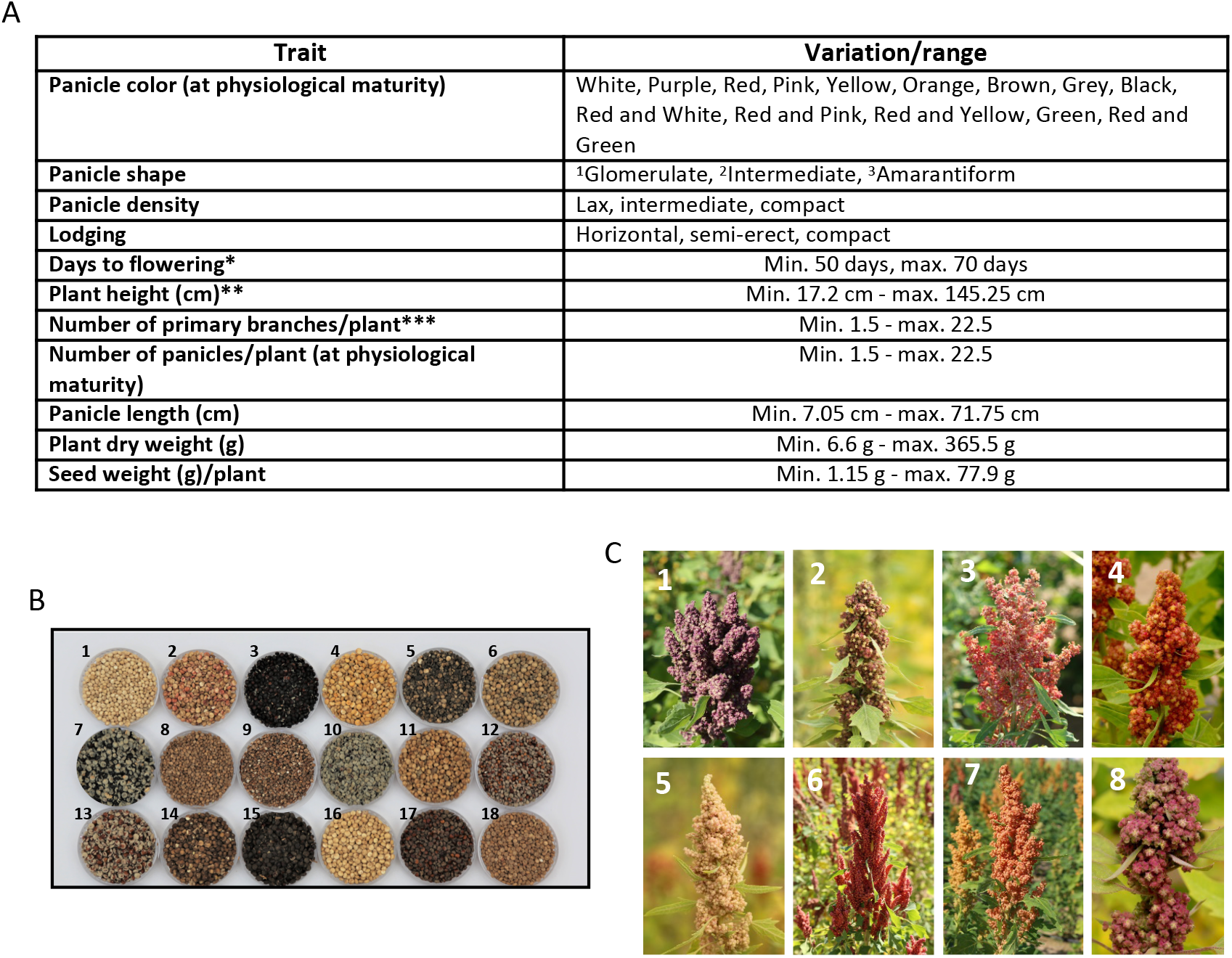
Morphological characteristics, and phenological and reproductive traits in quinoa genotypes. **(A)** Four qualitative and seven quantitative traits were measured and analyzed. ^1^Glomerulate: glomerules are inserted in the primary axis showing a globose shape; ^2^Intermediate: showing both shapes; ^3^Amarantiform: glomerules are inserted directly in the secondary axis and have an elongated shape. Asterisks indicate: * number of days from sowing until 50% of plants have started flowering; ** recorded at physiological maturity, from root collar to panicle apex; *** number of branches counted from the base to the second third of the stem from plants at physiological maturity. **(B)** Representative genotypes with diversity in seeds. **1.** CHEN 274, **2.** CHEN 210, **3.** D-12038, **4.** CHEN 109, **5.** D-9998, **6.** AMES 13219, **7.** D-12020, **8.** AMES 19046, **9.** PI-433378, **10.** D-12377, **11.** Co-Ka-1821, **12.** D-11999, **13.** CHEN 70, **14.** D-12140, **15.** CHEN 243, **16.** D-11980, **17.** D-12275, **18.** PI-433231. **(C)** Representative genotypes with diversity in panicle shape/color; **1.** Ames 13727, **2.** Ames 13749, **3.** PI 634924, **4.** Ames 13742, **5.** Ames 13761, **6.** Ames 22157, **7.** PI 665276, **8.** Ames 13731.

**Figure 2.**
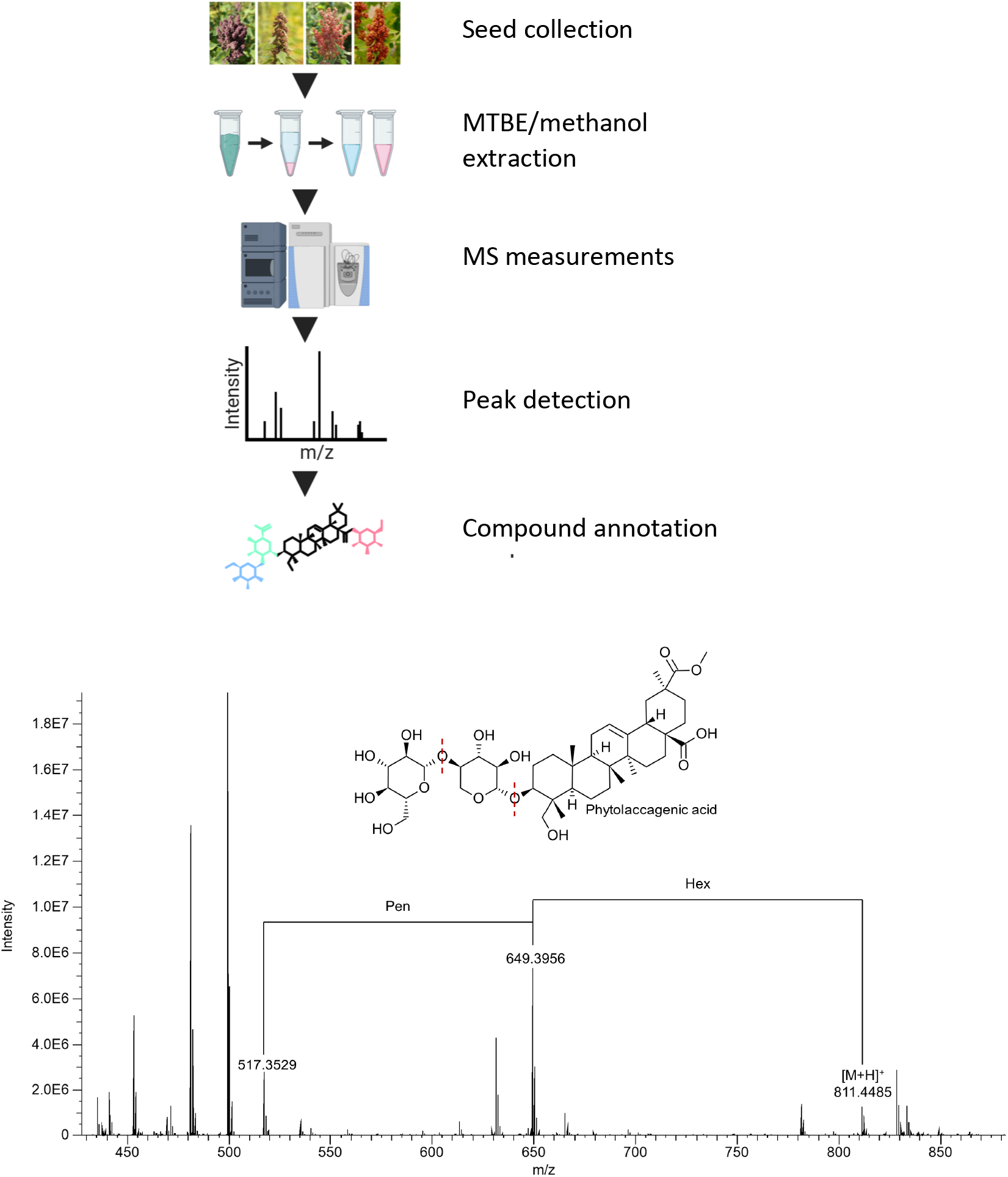
Schematic representation of the experimental pipeline. Seed samples were extracted using the MTBE/methanol method, followed by LC-MS analysis of the lipid and polar fractions, peak detection using GeneData software, and compound annotations using in-house reference libraries, and in-source fragmentation. The bottom panel displays an example of saponin annotation using in-source fragmentation in the positive mode. Given is a chromatogram and putative structure. The figure was prepared using BioRender (https://www.biorender.com).

**Figure 3.**
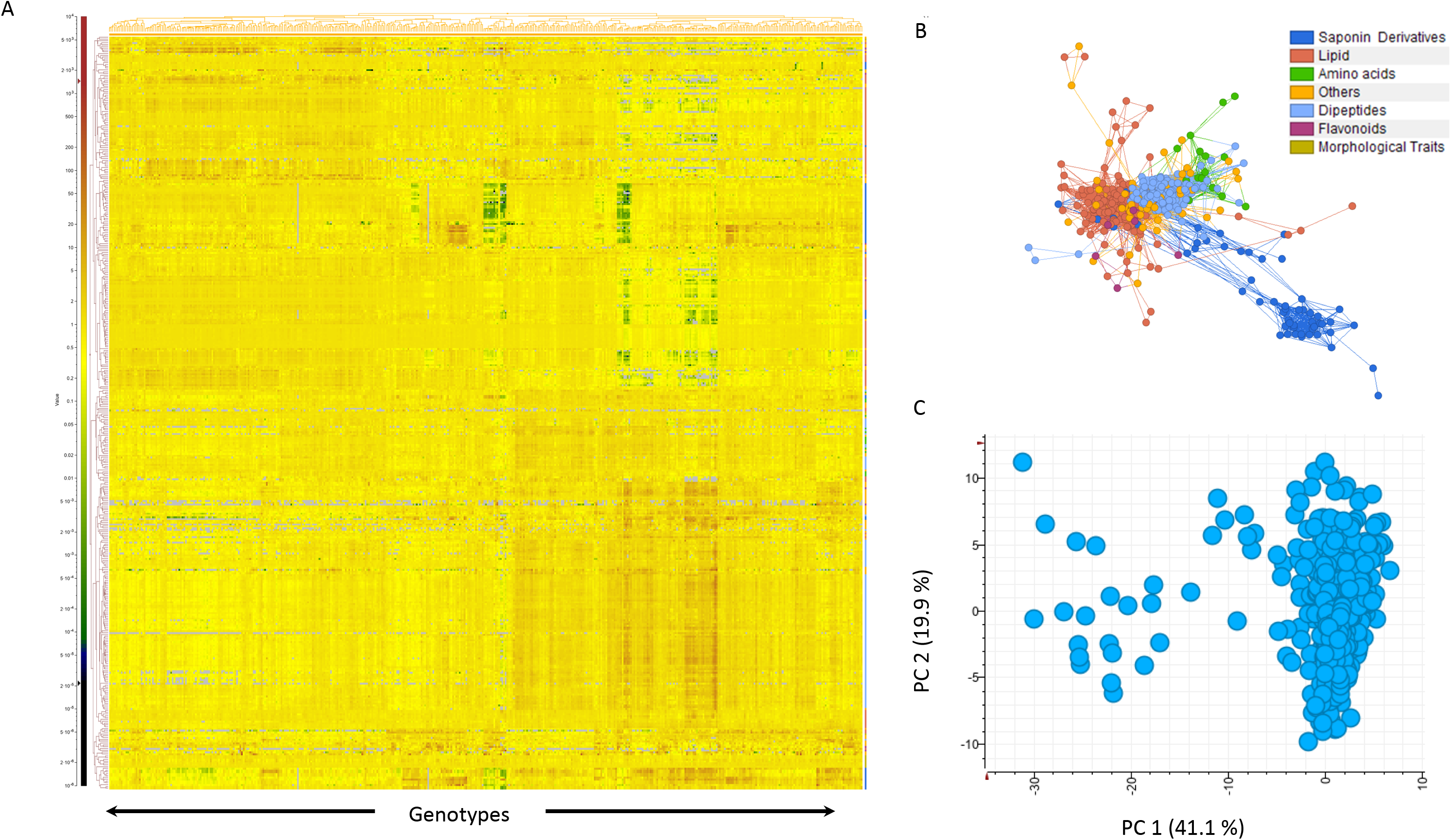
Phenotypic and metabolic variation across quinoa accessions. **(A)** Heatmap showing the relative levels of metabolites and phenotypic data measured on quinoa genotypes (full data set provided in **Supplementary Dataset 1** and **Supplementary Table 1**). **(B)** Correlation network between phenotypic traits and metabolic traits, each node represents a metabolite or a plant phenotypic trait, edges connecting two nodes show an association between two traits. **(C)** Principal component analysis (PCA) on saponin data across all genotypes.

## Methods

### Quinoa germplasm

The quinoa genotypes analyzed in this study (and their seed sources) are presented in **Supplementary Table 1**. Seeds of the different genotypes were propagated at the International Center for Biosaline Agriculture (ICBA) fields in years 2016 and 2017, and stored in a cold chamber at 2°C and a relative humidity of 30%.

### Experimental site and design

Experiments were carried out at the field research facilities of the International Center for Biosaline Agriculture, ICBA (N 25° 05.847; E 055° 23.464), Dubai, the United Arab Emirates, from November 2016 to April 2017. The soils at ICBA experimental fields are sandy in texture, that is, fine sand (sand 98%, silt 1%, and clay 1%), calcareous (50–60% CaCO_3_ equivalents), porous (45% porosity), and moderately alkaline (pH 8.2). The saturation percentage of the soil is 26 with a very high drainage capacity, while electrical conductivity of its saturation extract (ECe) is 1.2 dS m^−1^. According to the American Soil Taxonomy [20], the soil is classified as typic Torripsamments, carbonatic and hyperthermic [21]. Prior to sowing, poultry manure (Al Yahar Organic Fertilizers, UAE) was added at 40 tons per hectare (t ha^−1^) in the field chosen for the experiments. After four weeks of sowing, urea (nitrogen-phosphorus-potassium (NPK) content: 46-0-0) was applied at 40 kg ha^−1^, while NPK (20-20-20) was applied at 30 kg ha^−1^ after eight weeks of planting. Fertigation technique was used for the application of chemical fertilizers. The experimental plots were randomized following an augmented design [22], with each accession harboring a plot size of 1 m × 1 m. The distance between both, rows and plants was 25 cm.

### Irrigation system

A drip irrigation system was used for the experiment, with drippers at 25 cm distance, which was part of SCADA (Supervisory Control and Data Acquisition) system. Irrigation was provided twice a day for 10 min each time. For irrigation, about 13.3 L of water was used daily per plot. Data on relative humidity, temperature, and rainfall at the experimental site were recorded by the meteorological station at ICBA (**Supplementary Table 4**).

### Biological material

Four-hundred and sixty-eight quinoa genotypes, plus one accession from djulis (*Chenopodium formosanum* Koidz.) and one from goosefoot (*Chenopodium album* L.) were selected for the field experiment (**Supplementary Table 1**). Seeds from quinoa accession QQ74, for which a reference genome sequence is available [17], were included as well. The source of the seeds is given in **Supplementary Table 1**; seeds were kept in the active seed bank at ICBA, Dubai. The seeds were sown by hand by dibbling 2-3 seeds for each hole/location into the ground, to a depth of 1-2 cm near the dripper. Plants were thinned after about two weeks by removing unusually weak or strong individuals to leave one plant per location.

### Data collection

Eleven different morphological traits were recorded to assess the variation among the quinoa genotypes (including two species related to quinoa). For days to flowering, the data were recorded when about 50% of plants were flowering (Supplementary Table 1). Data on plant height, number of primary branches, number of panicles, main panicle length, plant dry weight, and seed weight were collected after plant maturity. For dry weight measurements, plants were kept in a drier electric oven (Model-PF 30, Carbolite, United Kingdom) at 40° for 48 hours.

### Extraction of lipids and polar metabolites

The extraction protocol was adapted and modified from Giavalisco et al. [23]. Metabolites were extracted from the quinoa seeds using a methyl-tert-butyl ether (MTBE)/methanol/water solvent system. Equal volumes of the lipid and polar fractions were dried in a centrifugal evaporator and stored at –20°C until processed further.

### LC-MS metabolomics

Polar and semipolar metabolites: After extraction, the dried aqueous phase was measured using ultra-performance liquid chromatography coupled to a Q-Exactive mass spectrometer (Thermo Fisher Scientific) in positive and negative ionization modes, as described [23]. Samples were run in ten consecutive sets of 50 samples and one set of 10 samples. Lipids: After extraction, the dried organic phase was measured using ultra-performance liquid chromatography coupled to a Q-Exactive mass spectrometer (Thermo Fisher Scientific) in positive mode, as described [23]. Samples were run in ten consecutive sets of 50 samples and one set of 10 samples.

### Data pre-processing: LC-MS metabolite data

Expressionist Refiner MS 12.0 (Genedata, Basel, Switzerland) was used for processing the LC-MS data (https://www.genedata.com/products/expressionist). Repetition was used to reduce the volume of data and to speed up processing. All types of data except Primary MS Centroid Data were removed using Data Sweep. Chemical Noise Subtraction activity was used to remove artefacts caused by chemical contamination. Snapshots of chromatograms were saved for further processing. Further processing of chromatogram snapshots was performed as follows: chromatogram alignment (Retention time (RT) search interval 0.5 min), peak detection (minimum peak size 0.03 min, gap/peak ratio 50%, smoothing window 5 points, centre computation by intensity-weighted method with intensity threshold at 70%, boundary determination using inflection points), isotope clustering (RT tolerance at 0.02 min, m/z tolerance 5 ppm, allowed charges 1 - 4), filtering for a single peak not assigned to an isotope cluster, charge and adduct grouping (RT tolerance 0.02 min, m/z tolerance 5 ppm). A detailed description of the software usage and possible settings was published before [24].

An MPI-MP in-house reference library was used to identify molecular features allowing 0.005 Da mass deviation and dynamic retention time deviation (maximum 0.2 min). Processing of fractionated samples resulted in annotation of 400 compounds (**Supplementary Table 2**).

Saponin and ecdysteroid annotation was based on the fragmentation behavior of the parent ion characteristic for the positive mode, and the mass of the main adduct measured in the negative mode.

### Data processing

Data represent normalised intensities of the main adduct measured in either the positive or negative mode. Normalisation was done to the median of a given metabolite calculated across a set.

### Data records

The raw metabolic data of the 471 samples were deposited to the MetaboLights database (https://www.ebi.ac.uk/metabolights/) [25] and are available under accession number MTBLS2382.

### Technical validation

To validate data reducibility, we chose 14 genotypes representing low and high saponin contents, and again analyzed the saponin content of their seeds (**Fig. 4; Supplementary Table 5**). The data showed high correlation (Pearson correlation coefficient, 0.98) between the sum of the saponin peaks identified in the two experiments validating metabolomics analysis.

**Figure 4.**
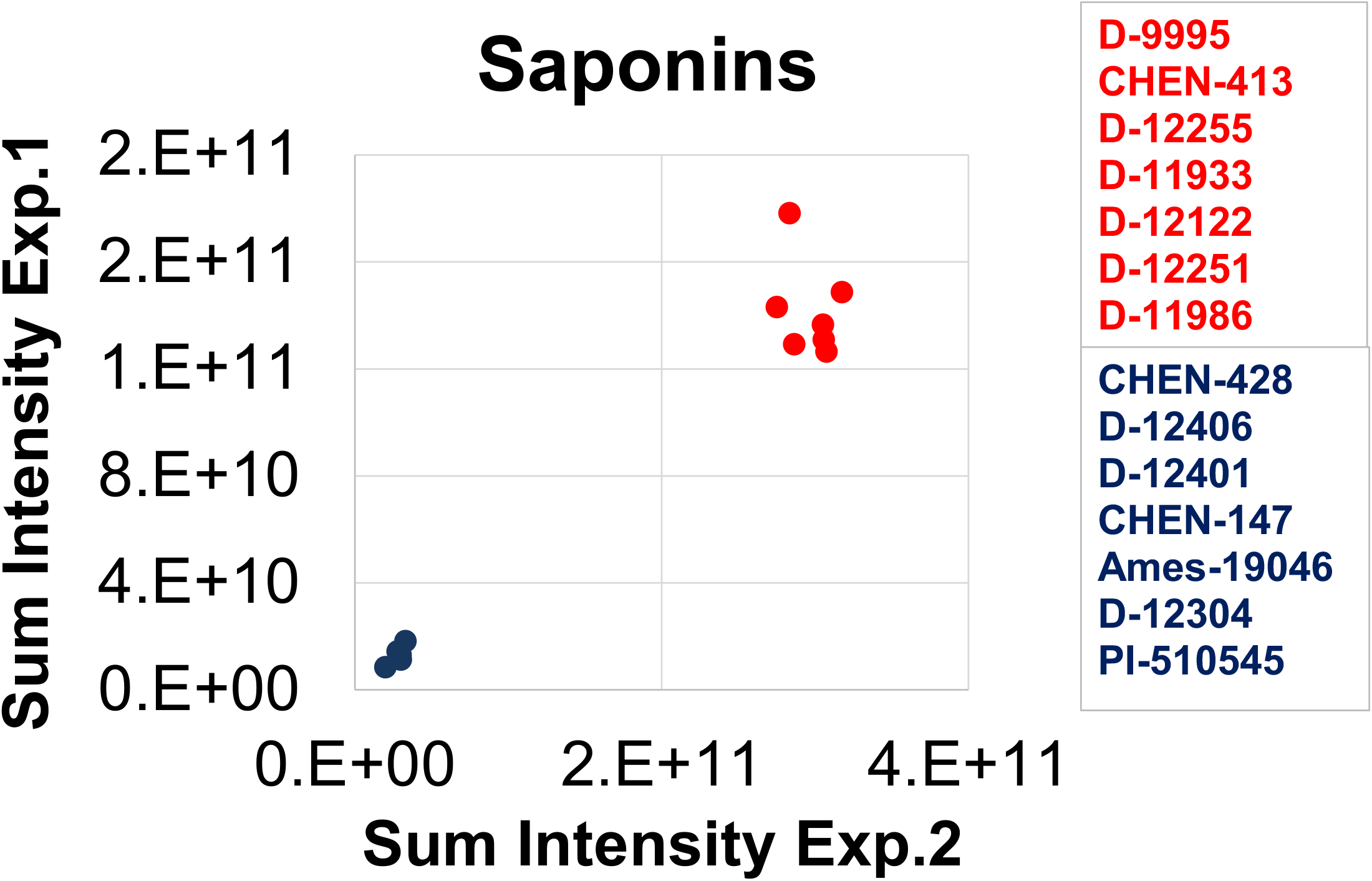
Seeds of 14 quinoa genotypes characterized by the lowest and highest saponin content. Metabolites were extracted and measured twice to assess metabolic profiling’s reproducibility. Data express a sum of all metabolic features detected in the positive mode in the retention time window between 8.14 and 14.00 min, which corresponds to the saponin elution and is used here as a proxy for total saponin content.

### Usage notes

In this study, we generated for the first time a large repertoire of the seed metabolome for 471 quinoa genotypes. Using high-resolution mass spectrometry, we annotated and provided normalized metabolite data of 400 compounds across the genotypes. Data are presented in EXCEL files (**Supplementary Dataset 1**). For each compound, we present m/z, retention time, ion detection mode, and annotation confidence (**Supplementary Table 2**) [23]. As mentioned above, the profiling data revealed considerable differences in metabolite abundances across genotypes. The data may be used to calculate the fold-change of certain metabolites between selected genotypes. In some cases, the composition of metabolites may influence product quality, as e.g. known for the Maillard reaction in bread making [26]. Hence, this dataset may be used in breeding programs when selecting specific genotypes with desirable metabolite profiles that may benefit product quality. Furthermore, in combination with the availability of genome sequences, the data can be used for functional genomics- and metabolite-based genome-wide association studies (mGWAS) to dissect the genetic basis of quinoa seed metabolism. The information on metabolite presence and quantity may also be used as a basis to design molecular markers to characterize responses to abiotic stresses. The data set is useful in genetic and correlation studies to investigate the relationship between metabolic diversity, geographical distribution, and integration with physiological and phenotypic diversity. The mass spectrometry raw data are available in MetaboLights, which allows download and re-processing with several commonly available tools such as xcm, GNPs and OpenMS. Furthermore, this enables the community to collect metabolite data of 471 different genotypes to generate a standard metabolome of quinoa.

### Code availability

Not applicable.

## Acknowledgements

S.B. thanks the Federal Ministry of Education and Research of Germany (BMBF) and the Arab-German Young Academy of Sciences and Humanities (AGYA) for funding of two Research Mobility Program grants to Dubai, United Arab Emirates, which allowed establishing the research reported here. B.M.-R. thanks the University of Potsdam, and S.B. thanks the Max Planck Institute of Molecular Plant Physiology for financial support. A.R.F. and B.M.-R. thank the European Union’s Horizon 2020 Research and Innovation Programme, project PlantaSYST (SGA-CSA No. 739582 under FPA No. 664620) for funding. All authors are very grateful to Rostyslav Braginets and Dirk Walther from the Max Planck Institute of Molecular Plant Physiology, Potsdam, for their great help in uploading the metabolomics data to the MetaboLights database.

## Author contributions

SB and BMR designed and coordinated the project. SB obtained funding. IT and EL extracted metabolites. AS analysed metabolite data. MW assisted with saponin annotation under the supervision of AS. SA performed PCA and clustering analysis under the supervision of ARF. MS, ST, and HM performed field experiments and collected the morphotrait data. KMM, SMS, and MT contributed to establish the seed collection at ICBA.

## Supplementary information

**Supplementary Table 1.** Morphological traits.

**Supplementary Table 2.** All detected peaks, putative metabolite names identified.

**Supplementary Table 3.** Genotypes exhibiting a correlation between their position in the PCA plot and their geographical origin.

**Supplementary Table 4.** Temperature, relative humidity, and rainfall at the trial site during the cropping season.

**Supplementary Table 5.** Seeds of 14 quinoa genotypes characterized by the lowest and highest saponin content.

**Supplementary Dataset 1.** All metabolite data.

