## Supplementary Table 4 for "The diversity of quinoa morphological traits and seed metabolic composition"

**Supplementary Table 4.** Temperature, relative humidity, and rainfall at the trial site during the cropping season.

| Year | Month | Temperature mean (℃) | Maximum temperature mean (℃) | Minimum temperature mean (℃) | Maximum relative humidity mean (%) | Minimum relative humidity mean (%) | Preci-pitation (mm) |
| --- | --- | --- | --- | --- | --- | --- | --- |
| 2016 | October | 28.3 | 35.9 | 20.7 | 74.9 | 20 | 0 |
| 2016 | November | 23.4 | 30.1 | 16.3 | 75.5 | 26.2 | 10.6 |
| 2016 | December | 20.3 | 26.8 | 13.7 | 80.0 | 29.8 | 14.6 |
| 2017 | January | 17.9 | 25.7 | 10.1 | 86.2 | 28.3 | 23.7 |
| 2017 | February | 24.2 | 32.4 | 16.1 | 77.3 | 22.1 | 17.9 |
| 2017 | March | 24.6 | 33.8 | 15.3 | 76.7 | 19.2 | 0 |
| 2017 | April | 27.2 | 35.8 | 21.4 | 71.9 | 17.7 | 0 |
| 2017 | May | 33.6 | 41.5 | 25.7 | 57.8 | 13.9 | 0 |
